# Synaptonemal complex components are required for meiotic checkpoint function in *C. elegans*

**DOI:** 10.1101/052977

**Authors:** Tisha Bohr, Guinevere Ashley, Evan Eggleston, Kyra Firestone, Needhi Bhalla

**Affiliations:** Department of Molecular, Cell and Developmental Biology, University of California, Santa Cruz, Santa Cruz, CA 95064

**Keywords:** meiosis, synapsis, chromosome, checkpoint, synaptonemal complex

## Abstract

Synapsis involves the assembly of a proteinaceous structure, the synaptonemal complex (SC), between paired homologous chromosomes and is essential for proper meiotic chromosome segregation. In *C. elegans*, the synapsis checkpoint selectively removes nuclei with unsynapsed chromosomes by inducing apoptosis. This checkpoint depends on Pairing Centers (PCs), *cis*-acting sites that promote pairing and synapsis. We have hypothesized that the stability of homolog pairing at PCs is monitored by this checkpoint. Here, we report that SC components SYP-3, HTP-3, HIM-3 and HTP-1 are required for a functional synapsis checkpoint. Mutation of these components does not abolish PC function, demonstrating they are bonafide checkpoint components. Further, we identify mutant backgrounds in which the instability of homolog pairing at PCs does not correlate with the synapsis checkpoint response. Altogether, these data suggest that, in addition to homolog pairing, SC assembly may be monitored by the synapsis checkpoint.

## Introduction

Meiosis is the specialized cell division by which cells undergo one round of DNA duplication and two successive rounds of division to produce haploid gametes from diploid organisms. During sexual reproduction, fertilization restores diploidy in the resulting embryo. In order for meiotic chromosomes to segregate properly in meiosis I and II, homologs pair, synapse and undergo crossover recombination (Bhalla *et al.* 2008). If homologous chromosomes fail to segregate properly, this can produce gametes, such as egg and sperm, with an improper number of chromosomes, termed aneuploidy. Embryos that result from fertilization of aneuploid gametes are generally inviable, but can also exhibit developmental disorders (Hassold and Hunt 2001). Therefore, checkpoint mechanisms monitor early meiotic prophase events to avoid the production of aneuploid gametes (MacQueen and Hochwagen 2011).

Synapsis involves the assembly of a proteinaceous complex, the synaptonemal complex (SC), between paired homologous chromosomes and is essential for crossover recombination (Bhalla and Dernburg 2008). In *C. elegans*, the synapsis checkpoint induces apoptosis to remove nuclei with unsynapsed chromosomes and prevent aneuploid gametes (Bhalla and Dernburg 2005) (Figure 1A). The synapsis checkpoint requires Pairing Centers (PCs) (Bhalla and Dernburg 2005), *cis*-acting sites near one end of each chromosome. PCs also promote pairing and synapsis (MacQueen *et al.* 2005) by recruiting factors, such as the zinc-finger containing proteins ZIM-1, ZIM-2, ZIM-3 and HIM-8 (Phillips and Dernburg 2006; Phillips *et al.* 2005), and the conserved polo-like kinase PLK-2 (Harper *et al.* 2011; Labella *et al.* 2011). We have hypothesized that the synapsis checkpoint monitors the stability of pairing at PCs as a proxy for proper synapsis (BOHR *et al.* 2015; Deshong *et al.* 2014). However, whether the process of synapsis is also monitored by the synapsis checkpoint is currently unknown.

**Figure 1.**
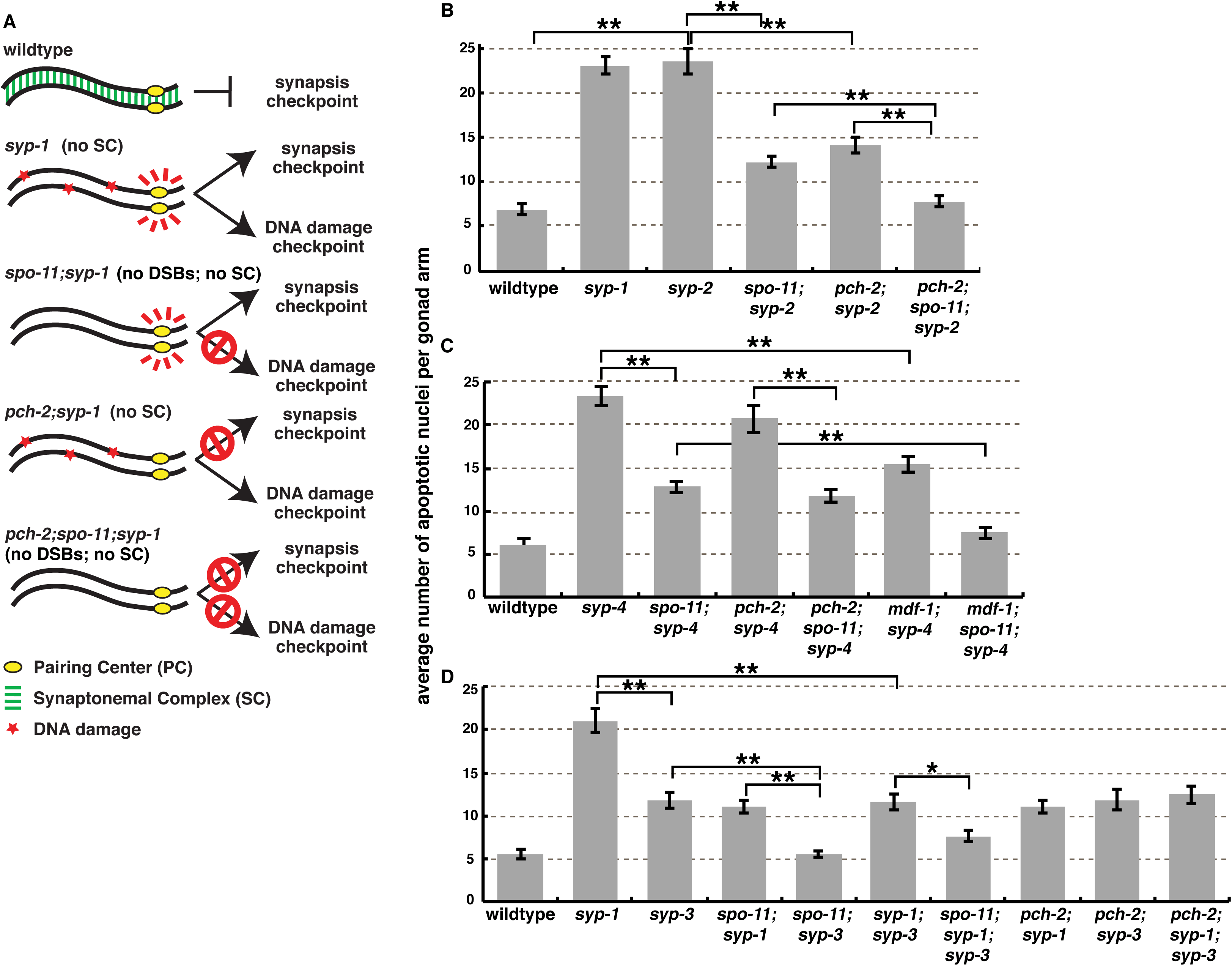
SYP-3 is required for the meiotic synapsis checkpoint. (A) Cartoons depicting meiotic checkpoint activation in *C. elegans*. (B) Elevation of germline apoptosis in *syp-2* mutants is dependent on *spo-11* and *pch-2*. (C) Elevation of germline apoptosis in *syp-4* mutants is dependent on *spo-11* and *mdf-1* but not on *pch-2*. (D) Elevation of germline apoptosis in *syp-3* mutants is dependent on *spo-11* but not on *pch-2*. Mutation of *syp-3* reduces apoptosis in *syp-1* and *syp-1;spo-11* double mutants but not *syp-1;pch-2* double mutants. Error bars represent ±SEM. A * indicates a p value < 0.01 and a ** indicates a p value < 0.0001.

Upon entry into meiosis, axial elements assemble between replicated sister chromatids to support homolog pairing and synapsis. In most species, HORMA domain proteins (HORMADs) associate with axial elements (Caryl *et al.* 2000; Fukuda *et al.* 2010; Hollingsworth *et al.* 1990; Wojtasz *et al.* 2009). These proteins share structural features with the well-characterized spindle checkpoint protein, Mad2 (Aravind and Koonin 1998; Kim *et al.* 2014), and have been implicated in monitoring meiotic prophase events, such as recombination and synapsis (Carballo *et al.* 2008; Daniel *et al.* 2011; Wojtasz *et al.* 2012), thus coupling meiotic chromosome architecture to checkpoint function. In *C. elegans*, four HORMAD proteins, HTP-3, HIM-3, HTP-1, and HTP-2, comprise the axial elements of the SC and play overlapping but distinct roles during meiotic prophase, including but not limited to meiotic checkpoint function (Couteau *et al.* 2004; Couteau and Zetka 2005; Goodyer *et al.* 2008; Kim *et al.* 2015; Martinez-perez and Villeneuve 2005; Zetka *et al.* 1999).

Synapsis is complete when the central element of the SC is assembled between paired axial elements of homologous chromosomes. In *C. elegans*, the central element includes the factors SYP-1, SYP-2, SYP-3 and SYP-4 (Colaiacovo *et al.* 2003; Macqueen *et al.* 2002; Smolikov *et al.* 2007; Smolikov *et al.* 2009). Loss of any one of these proteins produces a similar mutant phenotype: extensive asynapsis of all chromosomes that is accompanied by a delay in meiotic progression in which chromosomes remain asymmetrically localized in meiotic nuclei (Colaiacovo *et al.* 2003; MacQueen *et al.* 2002; Smolikov *et al.* 2007; Smolikov *et al.* 2009) and factors that normally localize transiently to meiotic chromosomes persist (Harper *et al.* 2011; Labella *et al.* 2011; Rosu *et al.* 2013; Stamper *et al.* 2013; Woglar *et al.* 2013). We’ve shown that *syp-1* mutants also induce germline apoptosis as a result of the synapsis checkpoint (Figure 1A) (Bhalla and Dernburg 2005). However, it’s unclear whether *syp-2*, *syp-3* or *syp-4* mutants similarly elicit an increase in germline apoptosis in response to the synapsis checkpoint. Genetically ablating the synapsis checkpoint does not affect the meiotic delay associated with asynapsis in *syp-1* mutants (Bohr *et al.* 2015; Deshong *et al.* 2014), indicating that these two events are not mechanistically coupled. Meiotic HORMAD proteins regulate this delay (Kim *et al.* 2015; Martinez-Perez and Villeneuve 2005).

Here, we report that some SC components are required for the synapsis checkpoint. *syp-2* mutants resemble *syp-1* mutants and elevate apoptosis in response to the synapsis checkpoint. *syp-4* mutants also exhibit elevated apoptosis similar to *syp-1* and *syp-2* mutants. However, the elevation in apoptosis observed in *syp-4* mutants is not dependent on PCH-2 but is dependent on MDF-1. Since both PCH-2 and MDF-1 are synapsis checkpoint components (Bhalla AND DERNBURG 2005; Bohr *et al.* 2015) that act redundantly to regulate synapsis (Bohr *et al.* 2015), these data suggest there may be molecular differences in how the synapsis checkpoint can be activated. By contrast, *syp-3* mutants do not elicit a synapsis checkpoint response, indicating that SYP-3 is required for the synapsis checkpoint. Similarly, *htp-3*, *him-3* and *htp-1* mutants are also defective in the synapsis checkpoint. The ability to generate a synapsis checkpoint response does not correlate with less stable homolog pairing at PCs, suggesting that the synapsis checkpoint may instead monitor SC assembly through these factors. Finally, loss of SYP-3, HTP-3, HIM-3 or HTP-1 does not abrogate PC function, consistent with these proteins playing a direct role in the checkpoint.

## Results and Discussion

### SYP-3 is required for the synapsis checkpoint

*syp-1* mutants exhibit increased germline apoptosis as a result of the synapsis checkpoint (due to asynapsis) and the DNA damage checkpoint (due to an inability to repair double strand breaks [DSBs]) (Figure 1A) (Bhalla and Dernburg 2005). SPO-11 is required for the introduction of meiotic DSBs (Dernburg *et al.* 1998) and PCH-2 is required for the synapsis checkpoint (Bhalla and Dernburg 2005). We’ve previously shown that loss of SPO-11 or PCH-2 in otherwise wild-type backgrounds does not affect germline apoptosis (Bhalla and Dernburg 2005). However, *spo-11;syp-1* and *pch-2;syp-1* double mutants display lower levels of germline apoptosis than *syp-1* single mutants because of loss of the DNA damage or synapsis checkpoint response, respectively. (Figure 1A) (Bhalla and Dernburg 2005). Loss of both checkpoints in *pch-2;spo-11;syp-1* triple mutants result in wild-type levels of apoptosis (Figure 1A) (Bhalla and Dernburg 2005).

To determine if other *syp* mutants behave similarly we quantified apoptosis in null *syp-2*, *syp-3* and *syp-4* mutants (Figure 1B, C and D). Mutation of *syp-2* elevated germline apoptosis levels similar to those seen in *syp-1* mutants (Figure 1B), suggesting that *syp-2* mutants exhibit both DNA damage and synapsis checkpoint responses. To verify that *syp-2* mutants exhibit a DNA damage checkpoint response, we introduced a mutation of *spo-11* into a *syp-2* background. We observed decreased apoptosis to intermediate levels in *spo-11;syp-2* double mutants (Figure 1B), indicating that *syp-2* mutants exhibit a DNA damage checkpoint response. To determine if *syp-2* mutants exhibit a synapsis checkpoint response we observed apoptosis in *pch-2;syp-2* double mutants which also had intermediate levels of germline apoptosis (Figure 1B). This verifies that *syp-2* mutants elevate germline apoptosis due to the synapsis checkpoint. Furthermore, mutation of both *pch-2* and *spo-11* reduced apoptosis to wild-type levels in a *syp-2* background (Figure 1B). These data show that the elevation of apoptosis observed in *syp-2* mutants is in response to both the DNA damage and synapsis checkpoints, similar to *syp-1* mutants (Bhalla and Dernburg 2005).

Next we analyzed *syp-4* mutants and found that germline apoptosis was also elevated (Figure 1C) comparable to *syp-1* and *syp-2* mutants (Figure 1B). Moreover, *spo-11;syp-4* double mutants resembled *spo-11;syp-1* and *spo-11;syp-2* double mutants (Bhalla and Dernburg 2005) (Figure 1B and C), demonstrating that *syp-4* mutants have elevated apoptosis due to the DNA damage checkpoint. However, germline apoptosis was unaffected in *pch-2;syp-4* and *pch-2;spo-11*; *syp-4* mutants compared to *syp-4* and *spo-11*; *syp-4* mutants, respectively (Figure 1C).

We reasoned that these results with the *pch-2* mutation could either reflect that an additional meiotic checkpoint was active in *syp-4* mutants or *syp-4* mutants produced a synapsis checkpoint response independent of PCH-2. We distinguished between these two possibilities by monitoring germline apoptosis in *mdf-1;syp-4* double mutants and *mdf-1;spo-11;syp-4;* triple mutants (Figure 1C). We previously reported that MDF-1, the *C. elegans* ortholog of the spindle checkpoint gene Mad1, is also required for the synapsis checkpoint and regulates synapsis in an independent, parallel pathway to PCH-2 (Bohr *et al.* 2015). Loss of MDF-1 reduced apoptosis to intermediate levels in *syp-4* mutants and wild-type levels in *spo-11;syp-4* mutants, indicating that the synapsis checkpoint contributes to the increase in apoptosis observed in *syp-4* mutants (Figure 1C). Thus, the genetic requirements for the synapsis checkpoint in *syp-4* mutants are different than that of *syp-1* and *syp-2* mutants.

We also quantified apoptosis in *syp-3* mutants and observed increased apoptosis compared to wild-type worms but not to levels observed in *syp-1* single mutants (Figure 1D). This suggests that unlike *syp-1, syp-2 and syp-4* mutants, *syp-3* mutants either have a functional DNA damage or synapsis checkpoint, but not both. To determine which checkpoint was responsible for the elevated apoptosis observed in *syp-3* mutants we first quantified apoptosis in *spo-11;syp-3* double mutants (Figure 1D). Mutation of *spo-11* in a *syp-3* background reduced apoptosis to wild-type levels (Figure 1D), demonstrating that the elevation in apoptosis observed in *syp-3* mutants is dependent on the DNA damage checkpoint. To ensure that the elevation in apoptosis observed in *syp-3* mutants is due solely to the DNA damage checkpoint and not due to the synapsis checkpoint, we monitored germline apoptosis in *pch-2;syp-3* mutants. Mutation of *pch-2* in the *syp-3* background did not reduce apoptosis (Figure 1D), illustrating that the elevation in apoptosis observed in *syp-3* mutants is not dependent on the synapsis checkpoint. Therefore, although chromosomes are unsynapsed in *syp-3* mutants (Smolikov *et al.* 2007), the synapsis checkpoint response is abrogated.

These data suggest that SYP-3 is required for the synapsis checkpoint. To verify this, we quantified apoptosis in *syp-1;syp-3* double mutants (Figure 1D). *syp-1;syp-3* double mutants had intermediate levels of germline apoptosis (Figure 1D), indicating loss of either the DNA damage checkpoint or the synapsis checkpoint but not both. Mutation of *syp-3* in a *pch-2;syp-1* background did not further decrease apoptosis (Figure 1D), confirming that SYP-3 is not required for the DNA damage checkpoint. However, *spo-11;syp-1;syp-3* triple mutants had wild-type levels of apoptosis (Figure 1D), signifying loss of the synapsis checkpoint. Altogether these data show that SYP-3, but not SYP-2 or SYP-4, is required for the synapsis checkpoint.

### *syp-3* and *syp-4* mutants exhibit more stable PC pairing than *syp-1* mutants

In the absence of synapsis (for example, in *syp-1* mutants), we can visualize pairing intermediates that typically precede and promote synapsis (MacQueen *et al.* 2002). Loss of PCH-2 further stabilizes pairing in *syp-1* mutants (Deshong *et al.* 2014), leading us to hypothesize that this stabilization of pairing, particularly at PCs, satisfies the synapsis checkpoint in *pch-2;syp-1* and *pch-2;syp-2* double mutants. We reasoned that since *syp-3* and *syp-4* mutants behaved differently than *syp-1* and *syp-2* mutants in the context of checkpoint activation, there might be similar differences with respect to PC pairing. We monitored pairing of *X* chromosomes as a function of meiotic progression by performing immunofluorescence against the PC protein HIM-8 (Phillips *et al.* 2005) in *syp-1*, *syp-3* and *syp-4* mutants, both in the presence and absence of PCH-2 (Figure 2A). Meiotic nuclei are arrayed in a spatiotemporal gradient in the germline, allowing for the analysis of the progression of meiotic events as a function of position in the germline (Figure 2B, see cartoon). We divided the germline into six equivalently sized zones and assessed the number of nuclei with paired HIM-8 signals in each zone. All six strains initiated pairing in zone 2, achieved maximal pairing by zone 4 and destabilized pairing in zones 5 and 6 (Figure 2B). Although we observed that loss of PCH-2 had effects on pairing in zone 6 in both *syp-3* and *syp-4* mutants (Figure 2B), signifying a role for PCH-2 in these backgrounds independent of the synapsis checkpoint, we focused our analysis on zone 2 based on the more stable pairing we detected in *pch-2;syp-1* double mutants in comparison to *syp-1* single mutants in this region (Figure 2B). PCs were more frequently paired in both *syp-3* and *syp-4* single mutants, similar to *pch-2;syp-1* mutants, in zone 2. *pch-2;syp-3* double mutants exhibited less steady-state pairing at *X* chromosome PCs than *syp-3* single mutants in zone 2, suggesting that in this background PCH-2 somehow promotes stable PC pairing. *pch-2;syp-4* double mutants resembled *syp-4* single mutants in zone 2, indicating that loss of PCH-2 in *syp-4* mutants does not further stabilize pairing at PCs and providing a potential explanation for why PCH-2 is not required for the synapsis checkpoint in *syp-4* mutants. Further, since *syp-4* mutants present similar frequencies of stable homolog pairing at PCs as *pch-2;syp-1* double mutants and nonetheless elicit a synapsis checkpoint response (Figure 1C) while *pch-2;syp-3* double mutants have paired PCs as infrequently as *syp-1* single mutants and do not activate germline apoptosis via the synapsis checkpoint (Figure 1D), these results suggest that stable PC pairing cannot be the sole criteria that satisfies the synapsis checkpoint.

**Figure 2.**
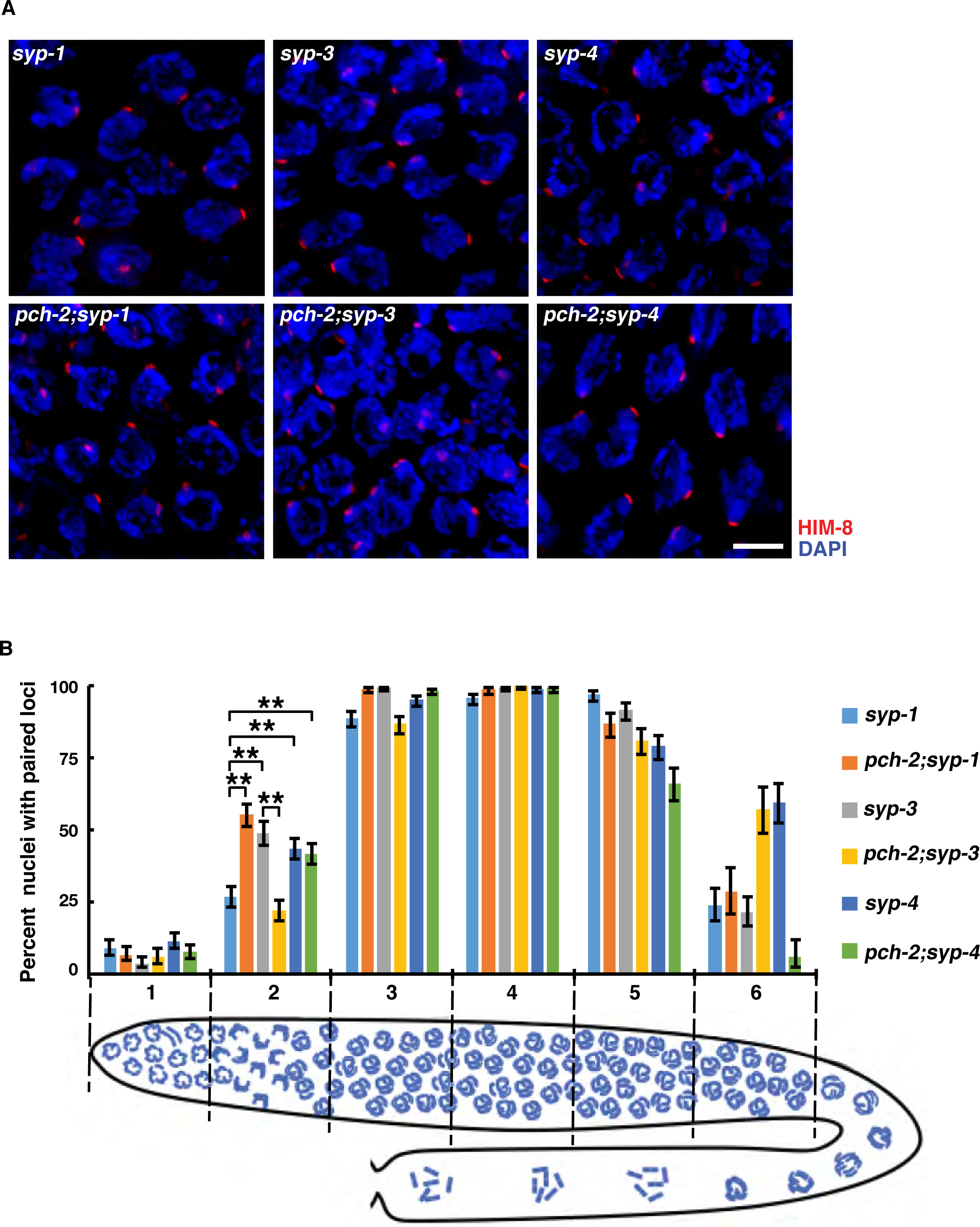
*syp-3* and *syp-4* mutants exhibit more stable PC pairing than *syp-1* mutants. (A) Images of meiotic nuclei in *syp-1*, *pch-2;syp-1*, *syp-3*, *pch-2;syp-3*, *syp-4* and *pch-2;syp4* mutants stained to visualize HIM-8 (red) and DNA (blue). Scale bars represent 5 µm. (B) Pairing at the *X* chromosome PC is more stable in *syp-3*, *syp-4* and *pch-2;syp-4* mutants than in *syp-1* mutants. The numbers on the x-axis correspond to regions of the gonad depicted in the cartoon in Figure 2B. Meiotic progression is from left to right. Error bars represent 95% confidence intervals. A ** indicates a p value < 0.0001. Significance was assessed by performing Fisher’s exact test.

### HORMAD proteins HTP-3, HIM-3 and HTP-1 are required for the synapsis checkpoint

We also tested whether axial element proteins, specifically HORMADs, are required for the 2 synapsis checkpoint using null mutations of each gene (Figure 3). First, we tested whether HTP-3 and HIM-3 are required for the synapsis checkpoint by monitoring apoptosis in *htp-3* and *him-3* mutants (Figure 3A). *htp-3* and *him-3* mutants produced wild-type levels of apoptosis (Figure 3A), despite their inability to synapse chromosomes (Couteau *et al.* 2004; Goodyer *et al.* 2008; Zetka *et al.* 1999). Thus, these mutants produce neither a DNA damage checkpoint nor a synapsis checkpoint response. HTP-3 is required for DSB formation in meiosis (Goodyer *et al.* 2008) and HIM-3 is thought to promote inter-homolog recombination by inhibiting inter-sister repair (Couteau *et al.* 2004). These phenotypes could explain the inability of these mutants to generate a DNA damage response. To further investigate a possible role for HTP-3 and HIM-3 in the synapsis checkpoint, we introduced mutations of *htp-3* and *him-3* into *syp-1* mutants and quantified apoptosis. *syp-1;htp-3* and *syp-1;him-3* double mutants have wild-type levels of germline apoptosis (Figure 3A), demonstrating that, even in the *syp-1* background, HTP-3 and HIM-3 are indeed required for the synapsis checkpoint.

**Figure 3.**
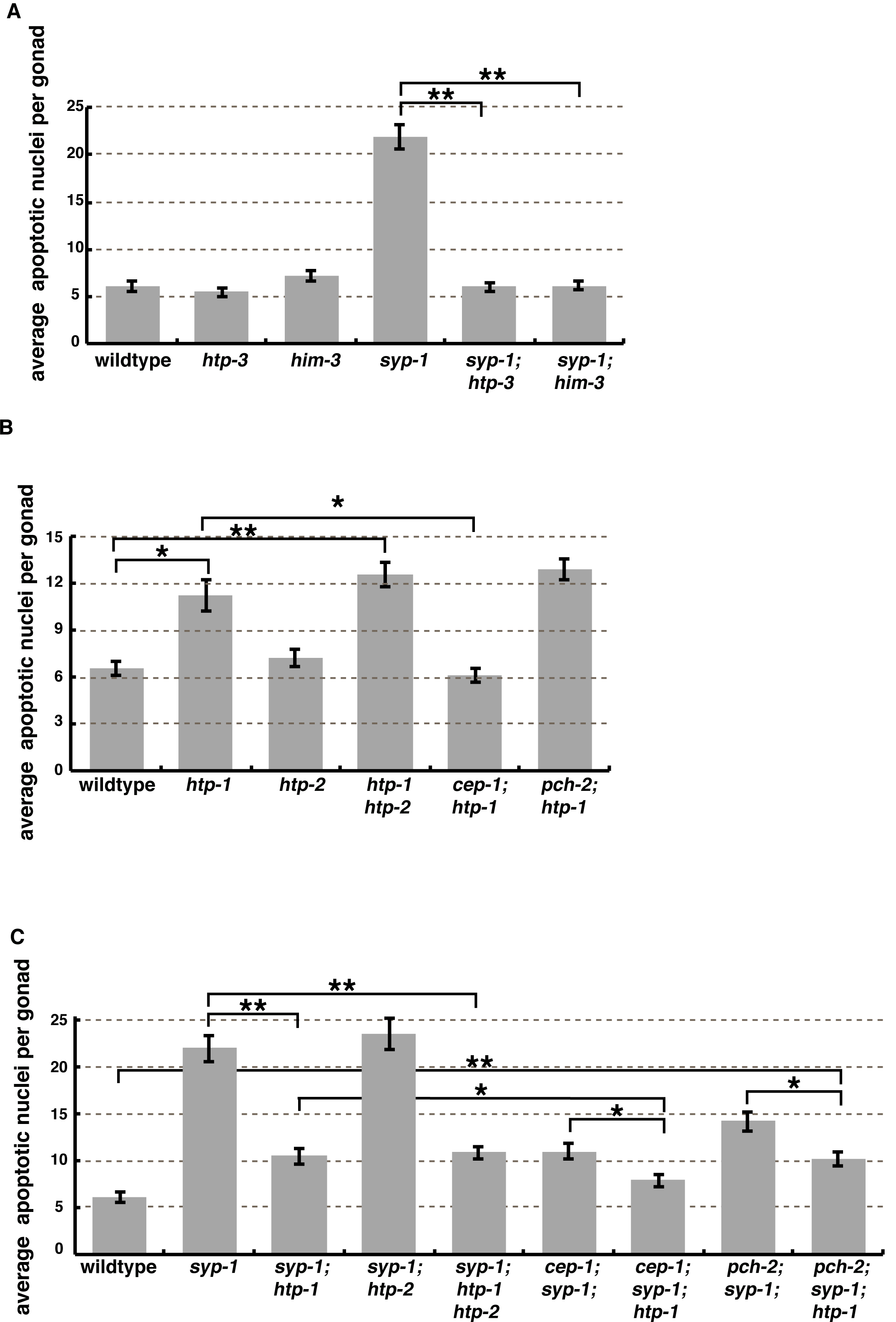
HTP-3, HIM-3 and HTP-1 are required for the synapsis checkpoint. (A) *htp-3 and him-3* mutants have wild-type levels of germline apoptosis and reduce germline apoptosis in *syp-1* mutants. (B) The elevation of germline apoptosis in *htp-1* mutants is *cep-1* dependent but not *pch-2* dependent. (C) Mutation of *htp-*1 reduces germline apoptosis in *syp-1* single and *cep-1;syp-1* double mutants. Error bars represent ±SEM. A * indicates a p value < 0.01 and a ** indicates a p value < 0.0001.

We then tested whether HTP-1 and HTP-2 are required for the synapsis checkpoint. *htp-1* single mutants synapse their chromosomes non-homologously (Couteau and Zetka 2005; Martinez-perez and Villeneuve 2005) and had intermediate levels of apoptosis (Figure 3B). These data suggest that *htp-1* mutants elicit a DNA damage or synapsis checkpoint response but not both. *htp-2* single mutants have no obvious meiotic defects (Couteau and Zetka 2005) and exhibited wild-type levels of apoptosis (Figure 3B), indicating that *htp-2* mutants do not produce a DNA damage or synapsis checkpoint response. *htp-1* is linked to *spo-11* on chromosome IV, making it difficult to create *spo-11 htp-1* double mutants. Therefore, to investigate which checkpoint was responsible for the intermediate levels of apoptosis observed in *htp-1* mutants, we abrogated the DNA damage checkpoint using a mutation in *cep-1*, the *C. elegans p53* orthologue (Derry *et al.* 2001; Schumacher *et al.* 2001). Mutation of *cep-1* in the *htp-1* background reduced apoptosis to wild-type levels while mutations of *pch-2* had no effect on germline apoptosis when compared to *htp-1* single mutants (Figure 3B). This indicates that the elevation in apoptosis observed in *htp-1* mutants is dependent on the DNA damage checkpoint and not the synapsis checkpoint. Therefore, unlike *htp-3* and *him-3* mutants (Figure 3A), *htp-1* mutants activate germline apoptosis in response to the DNA damage checkpoint (Figure 3B), supporting the idea that meiotic HORMADS also play distinct roles during meiotic checkpoint activation. Furthermore, these data suggest that either non-homologous synapsis does not result in a synapsis checkpoint response or that HTP-1 may be required for the synapsis checkpoint.

To test if HTP-1 is required for the synapsis checkpoint, we took advantage of the partially redundant roles of HTP-1 and HTP-2 during meiotic synapsis. *htp-1 htp-2* double mutants have unsynapsed chromosomes (Couteau and Zetka 2005), similar to *htp-3* and *him-3* single mutants (Couteau *et al.* 2004; Goodyer *et al.* 2008; Zetka *et al.* 1999), allowing us to evaluate whether unsynapsed chromosomes elicit a synapsis checkpoint response in the absence of HTP-1. Similar to *htp-1* single mutants, *htp-1 htp-2* double mutants exhibited intermediate apoptosis (Figure 3B), suggesting that abrogation of the synapsis checkpoint in *htp-1* mutants is not the product of non-homologous synapsis and supporting the possibility that HTP-1 is required for the synapsis checkpoint. In addition, these data demonstrate that HTP-1 and HTP-2 do not appear to play redundant roles in the DNA damage checkpoint’s induction of germline apoptosis. This is in contrast to the redundant roles they play in regulating meiotic progression when chromosomes are unsynapsed (Kim *et al.* 2015).

To validate that HTP-1 is required for the synapsis checkpoint we observed apoptosis in *syp-1;htp-1* and *syp-1;htp-2* double mutants (Figure 3C). While mutation of *htp-2* had no effect on apoptosis in the *syp-1* background, we observed reduced apoptosis to intermediate levels in *syp-1;htp-1* double mutants compared to *syp-1* single mutants (Figure 3C), indicating loss of one checkpoint. To verify that the synapsis checkpoint but not the DNA damage checkpoint is abrogated in the *syp-1;htp-1* background we observed apoptosis in *pch-2*;*syp-1;htp-1 and cep-1;syp-1*; *htp-1* triple mutants. Mutation of *cep-1* in the *syp-1;htp-1* background reduced apoptosis to levels comparable to wild-type worms (Figure 3C) demonstrating that the elevation of apoptosis observed in *syp-1*; *htp-1* mutants is dependent on the DNA damage checkpoint. In addition, mutation of *pch-2* did not further decrease apoptosis in the *syp-1*; *htp-1* background (Figure 3C), showing that the elevation of apoptosis observed in *syp-1*; *htp-1* mutants is not dependent on the synapsis checkpoint. Therefore, the synapsis checkpoint is abrogated in *syp-1*; *htp-1* mutants. However, while apoptosis in *pch-2*; *syp-1;htp-1* triple mutants was significantly higher than wild-type, *pch-2*; *syp-1;htp-1* triple mutants had reduced levels of apoptosis in comparison to *pch-2;syp-1* double mutants (Figure 3C), suggesting that loss of HTP-1 affects the synapsis checkpoint more severely than loss of PCH-2. Alternatively, loss of HTP-1 may partially reduce the DNA damage response in this background given its role in enforcing meiotic-specific DNA repair mechanisms (Martinez-perez and Villeneuve 2005). Lastly, similar to *syp-1;htp-1* double mutants, *syp-1;htp-1 htp-2* triple mutants exhibited intermediate levels of apoptosis compared to *syp-1* single mutants and wild-type worms (Figure 3C), further verifying that HTP-2 is not redundant with HTP-1 when considering checkpoint activation of apoptosis. Altogether, these data illustrate that HTP-3, HIM-3, and HTP-1, but not HTP-2, are required for the synapsis checkpoint.

### HTP-3 and HIM-3 disrupt localization of some but not all PC proteins

HTP-3, HIM-3 and HTP-1 could be directly required for the synapsis checkpoint or these proteins could be involved in regulating other mechanisms that are required for the synapsis checkpoint. For example, since PCs are required for the synapsis checkpoint (Bhalla and Dernburg 2005), we were concerned that *htp-3, him-3* and *htp-1* mutants might have defects in PC function. Since *htp-1* single mutants produce non-homologous synapsis (Couteau and Zetka 2005; Martinez-perez and Villeneuve 2005) and our analysis of apoptosis shows that loss of HTP-2 has no effect on synapsis checkpoint signaling (Figures 3C), we performed experiments to address this using *htp-1 htp-2* double mutants, which have unsynapsed chromosomes (Couteau and Zetka 2005), allowing better comparison with *htp-3* and *him-3* single mutants. We localized ZIM-2, a protein that binds to and is required for PC function of Chromosome V (Phillips and Dernburg 2006), in wild-type worms and *htp-3*, *him-3* and *htp-1 htp-2* mutants in early meiotic prophase nuclei (Figure 4A). In wild-type worms ZIM-2 forms robust patches at the nuclear periphery (Figure 4A) (Phillips and Dernburg 2006). We observed ZIM-2 staining in *htp-1 htp-2* double mutants similar to wild-type worms (Figure 4A). However, *htp-3* and *him-3* mutants had less robust ZIM-2 localization compared to wild-type worms (Figure 4A). We saw similar results in *htp-3*, *him-3* and *htp-1 htp-2* mutants when we stained for ZIM-1 and ZIM-3 (Figure S1A and B), which bind the PCs of Chromosomes I and IV and Chromosomes II and III, respectively (Phillips and Dernburg 2006).

**Figure 4:**
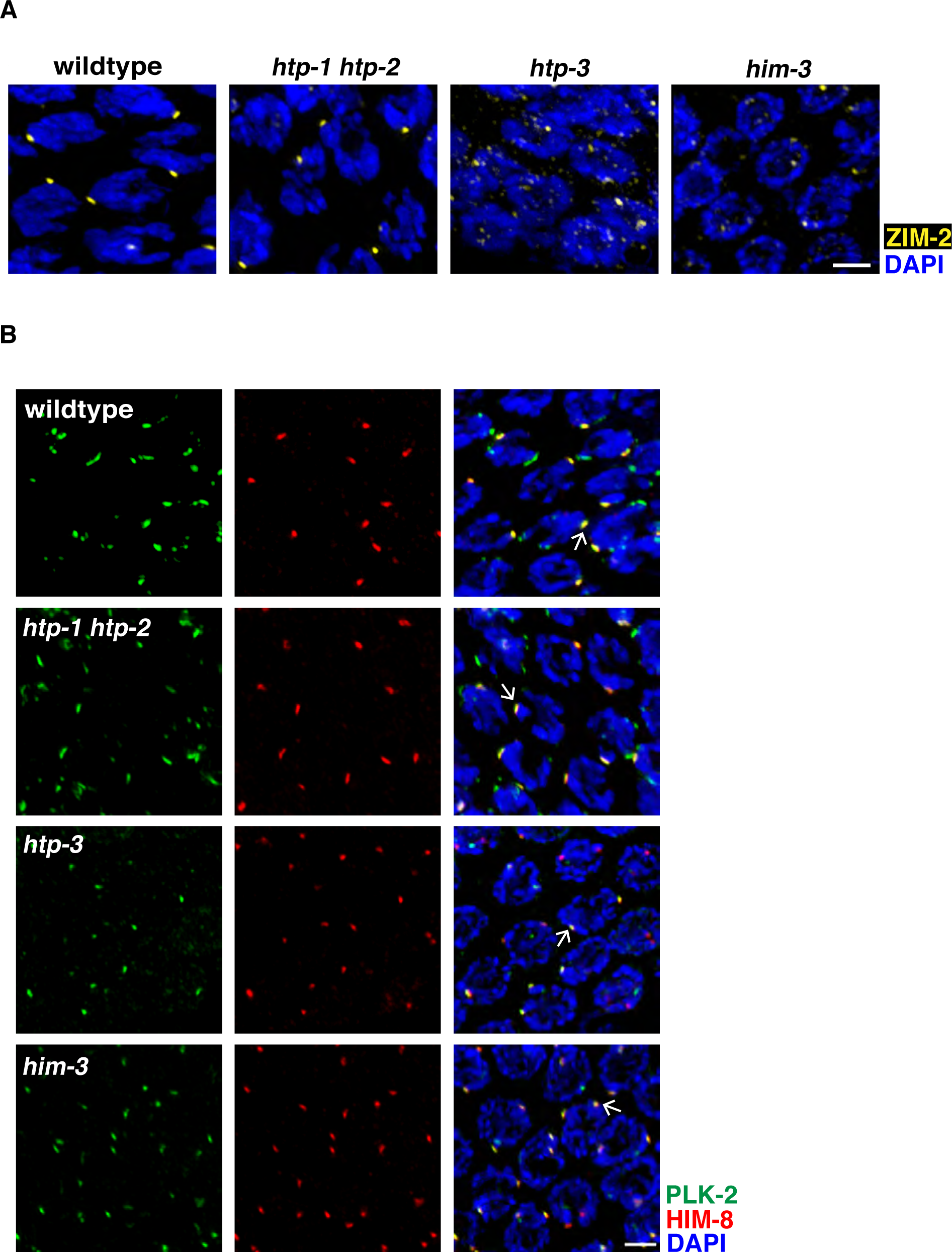
Loss of HTP-3 and HIM-3 disrupts localization of some but not all PC proteins. (A) Images of early meiotic prophase nuclei in wild-type worms, *htp-1/2, htp-3,* and *, him-3* mutants stained to visualize ZIM-2 (yellow) and DAPI (blue). (B) Images of early meiotic prophase nuclei in wild-type worms, *htp-1/2, htp-3,* and *, him-3* mutants stained to visualize PLK-2 (green), HIM-8 (red) and DAPI (blue). Arrow indicates an example of colocalization of PLK-2 and HIM-8. Scale bar represents 2 µm.

The defect in robustly localizing ZIMs to PCs in *htp-3* and *him-3* mutants (Figure 4A, S1A and B) might explain why these mutants are defective in the synapsis checkpoint. However, a single unsynapsed *X* chromosome, with an active PC, is sufficient to elicit a checkpoint response (Bhalla and Dernburg 2005). Therefore, we also localized the *X* chromosome PC binding protein, HIM-8 (Figure 4B) (Phillips *et al.* 2005). We observed staining patterns similar to wild-type worms in *htp-3*, *him-3* and *htp-1 htp-2* mutants (Figure 4B). However, consistent with published reports (Couteau *et al.* 2004; Couteau and Zetka 2005; Goodyer *et al.* 2008), HIM-8 foci were more often unpaired in *htp-3* and *him-3* mutants, while in wild-type and *htp-1 htp-2* double mutants a single HIM-8 focus per nucleus could often be observed in early meiotic prophase nuclei. We also determined whether *X* chromosome PCs were functional in these mutant backgrounds by localizing PLK-2 (Figure 4B), a kinase that is recruited by PCs to promote synapsis and the synapsis checkpoint (Harper *et al.* 2011; Labella *et al.* 2011). In *htp-3*, *him-3* and *htp-1 htp-2* mutants, PLK-2 co-localized with HIM-8 (Figure 4B), indicating *X* chromosome PCs were active. Altogether, these data argue against the interpretation that mutations in HORMAD proteins abrogate the synapsis checkpoint indirectly due to defects in PC function and support the conclusion that they are involved in the synapsis checkpoint response.

### *syp-3* mutants have active PCs

Similar to *htp-3*, *him-3* and *htp-1 htp-2* mutants, *syp-3* mutants have unsynapsed chromosomes but fail to elevate germline apoptosis in response to the synapsis checkpoint (Figure 1D). Unlike *htp-3*, *him-3* and *htp-1 htp-2* mutants, *syp-3* mutants display a delay in meiotic progression (Smolikov *et al.* 2007), likely because HTP-3, HIM-3, HTP-1 and HTP-2 are present to promote this delay (Kim *et al.* 2015; Martinez-perez and Villeneuve 2005). However, this delay in meiotic progression does not depend on PC function (Kim *et al.* 2015), raising the possibility that *syp-3* mutants abrogate the synapsis checkpoint due to defective PCs. To directly test this, we localized PLK-2 in meiotic prophase in *syp-3* mutants and compared them to wild-type worms, *syp-1*, *syp-2* and *syp-4* mutants. Similar to wild-type animals and *syp-1* (Harper *et al.* 2011; Labella *et al.* 2011), *syp-2* and *syp-4* mutants, *syp-3* mutants robustly localized PLK-2 to PCs (Figure 5A). Moreover, unlike wild-type germlines, PLK-2 localization on PCs is extended in *syp-3* mutants, similar to *syp-1*, *syp-2* and *syp-4* mutants (Figure 5A).

**Figure 5:**
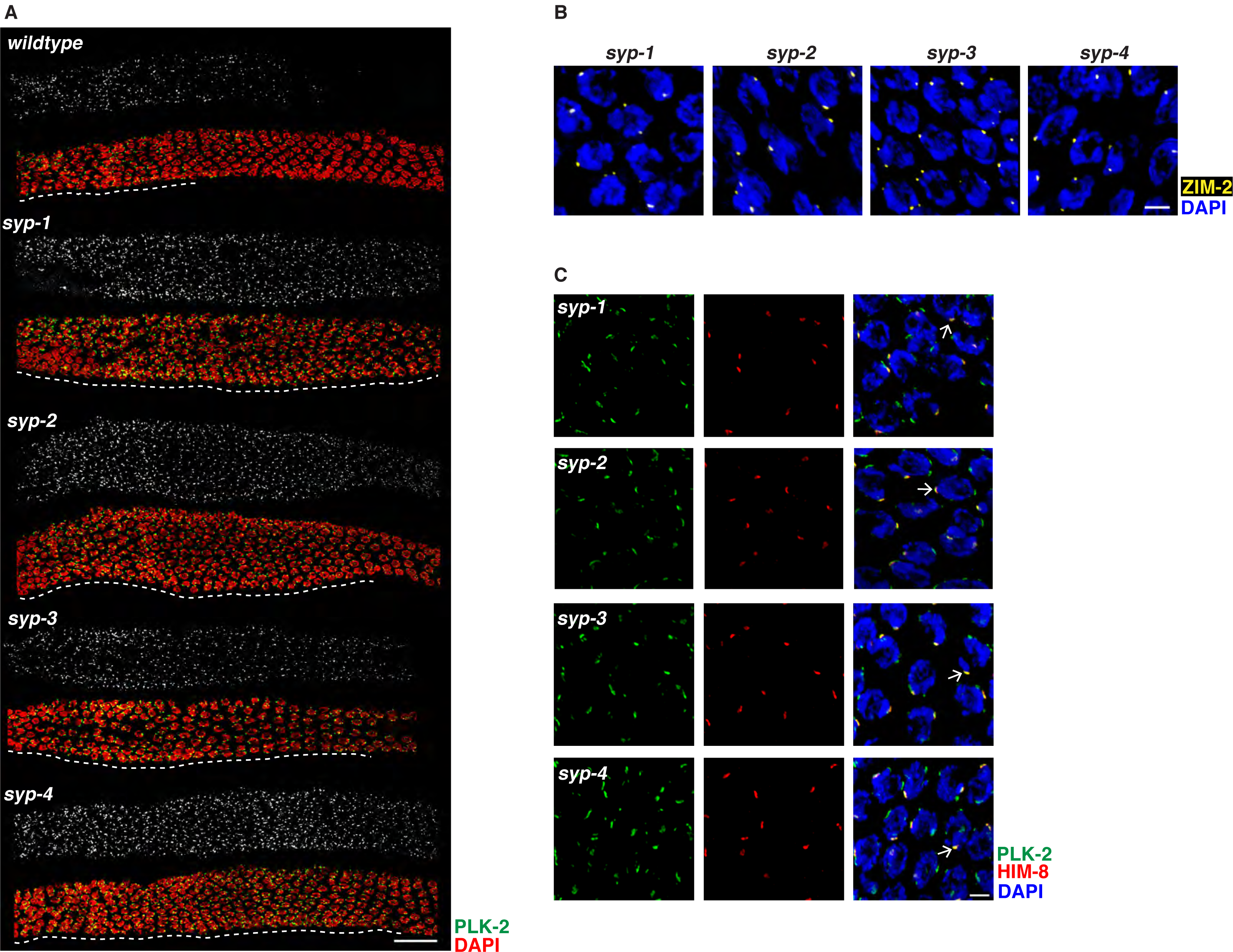
*syp-3* mutants have active PCs. (A) Images of germlines, from entry into meiosis until late meiotic prophase, of wild-type worms, *syp-1*, *syp-2, syp-3*, and *syp-4* mutants stained to visualize PLK-2 (green and grayscale) and DAPI (red). Delay in meiotic progression indicated by white dashed line. Scale bar represents 30 µm. (B) Images of early meiotic prophase nuclei in wild-type worms, *syp-1*, *syp-2, syp-3*, and *syp-4* mutants stained to visualize ZIM-2 (yellow) and DAPI(blue). (C) Images of early meiotic prophase nuclei in wild-type worms, *syp-1*, *syp-2, syp-3*, and *syp-4* mutants stained to visualize PLK-2 (green), HIM-8 (red) and DAPI (blue). Arrow indicates an example of colocalization of PLK-2 and HIM-8. Scale bar represents 2 µm.

We complemented this evaluation of PC function by localizing ZIM-2 and HIM-8 in *syp-3* mutants and compared this to *syp-1*, *syp-2* and *syp-4* mutants. ZIM-2 forms robust patches in meiotic nuclei in *syp-3* mutants, similar to *syp-1, 2 and 4* mutants (Figure 5B). Furthermore, HIM-8 localizes to all meiotic nuclei in *syp-3* mutants and co-localizes with PLK-2 (Figure 5C). These data show that SYP-3 is required for the synapsis checkpoint in a mechanism distinct from regulating PC function.

Altogether, our data demonstrate that some SC components, namely SYP-3, HTP-3, HIM-3 and HTP-1, are required for the synapsis checkpoint (Figure 6). Furthermore, their involvement in the synapsis checkpoint does not correlate with their effects on PC pairing (Figure 2 and Couteau *et al.* 2004; Couteau AND ZETKA 2005; Goodyer *et al.* 2008), suggesting they contribute to synapsis checkpoint function in some unique fashion. We propose that the synapsis checkpoint monitors SC assembly via these SC components. Uncovering which specific functions of SYP-3 and the HORMADs are required for the synapsis checkpoint are intriguing questions to be addressed in future studies.

**Figure 6:**
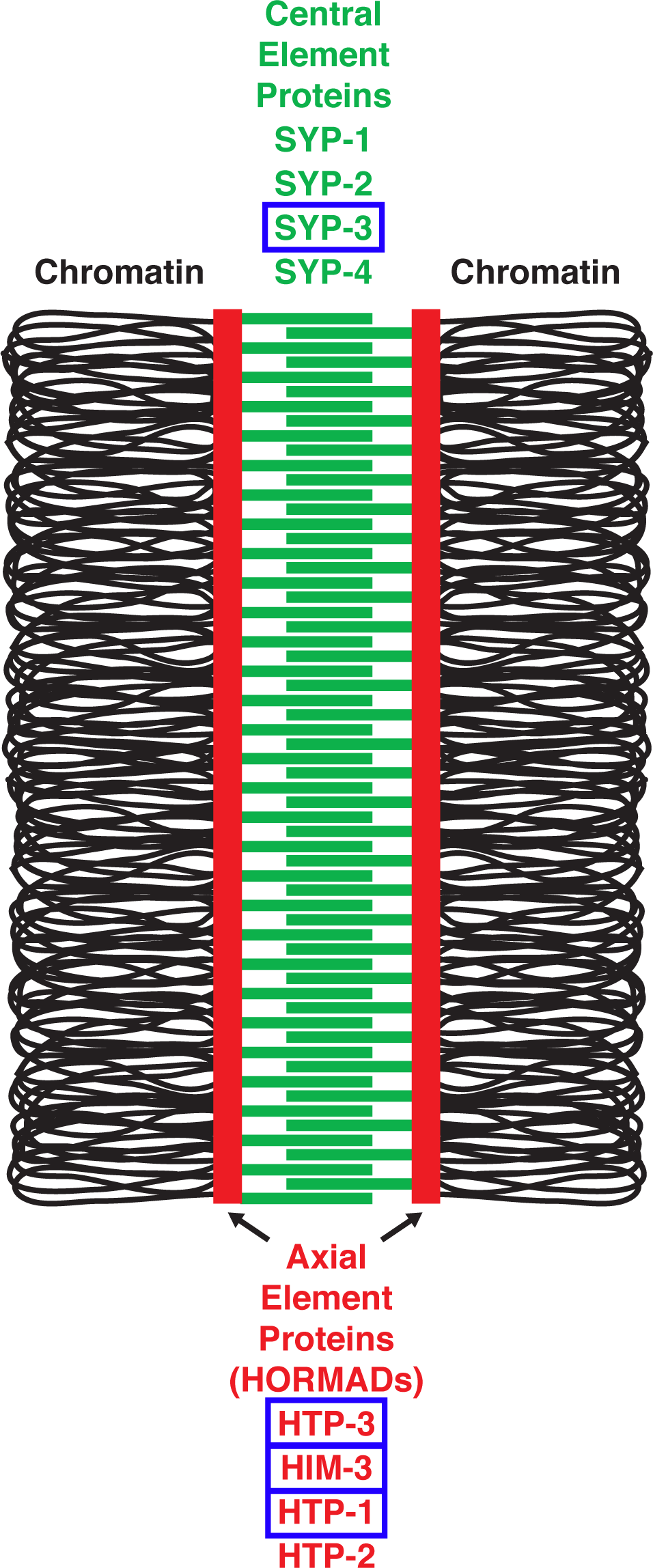
Cartoon of the synaptonemal complex (SC) in *C. elegans*. Central element components are in green (SYP-1, SYP-2, SYP-3 and SYP-4) and axial element components (HORMADs) are in red (HTP-3, HIM-3, HTP-1 and HTP-2). Chromatin is depicted as black loops tethered by axial elements. SC components that are required for the synapsis checkpoint are boxed in blue.

Surprisingly, despite having similar defects in synapsis, we found that not all central element components of the SC are equivalent in the context of checkpoint function. While *syp-2* mutants essentially phenocopy *syp-1* mutants, *syp-4* mutants have a functional synapsis checkpoint that is independent of PCH-2 but dependent on MDF-1. When combined with our pairing analysis (Figure 2B), these data raise the possibility that SYP-4 could be playing another role during the synapsis checkpoint. SYP-4 was identified by virtue of its two-hybrid interaction with SYP-3. However, unlike SYP-3, SYP-4 does not show an interaction with either SYP-1 or SYP-2 by two-hybrid (Smolikov *et al.* 2009). While there are a variety of reasons why relevant protein-protein interactions might not be recapitulated by yeast two-hybrid assays, these negative data suggest that SYP-4 could uniquely interact with SYP-3 during synapsis. For example, one scenario consistent with our data is that when SYP-3 is not bound to SYP-4, SYP-3 signals to the synapsis checkpoint and when it is bound to SYP-4, this signal is silenced. Future experiments will address this hypothesis.

## Materials and Methods

### Genetics and Worm Strains

The wildtype *C. elegans* strain background was Bristol N2 (Brenner 1974). All experiments were performed on adult hermaphrodites at 20° under standard conditions. Mutations and rearrangements used were as follows:

> LG I: *htp-3(tm3655)*, *syp-4 (tm2713)*, *cep-1(gk138)*, *syp-3(ok258)*, *hT2 [bli-4(e937) let-?(q782) qIs48]* (I;III)
>
> LG II: *pch-2(tm1458)*
>
> LG IV: *htp-1(gk174)*, *htp-2(tm2543)*, *him-3(gk149)*, *spo-11(ok79)*, *nT1[unc-?(n754) let-?(m435)] (IV, V)*, *nT1 [qIs51]*(IV, V)
>
> LG V: *syp-2(ok307), syp-1(me17), mdf-1(av19), bcIs39(Pim::ced-1::GFP)*

### Quantification of Germline Apoptosis

Scoring of germline apoptosis was performed as previously descried in (Bhalla and Dernburg 2005). L4 hermaphrodites were allowed to age for 22 hours at 20°C. Live worms were mounted under coverslips on 1.5% agarose pads containing 0.2mM levamisole. A minimum of twenty-five germlines were analyzed for each genotype by performing live fluorescence microscopy and counting the number of cells fully surrounded by CED-1::GFP. Significance was assessed using a paired t-test between all mutant combinations. All experiments were performed at least twice.

### Antibodies, Immunostaining and Microscopy

Immunostaining was performed on worms 20 to 24 hours post L4 stage. Gonad dissection were carried out in 1X EBT (250 mM HEPES-Cl pH 7.4, 1.18 M NaCl, 480 mM KCl, 20 mM EDTA, 5 mM EGTA) + .1% Tween 20 and 20mM sodium azide. An equal volume of 2% formaldehyde in EBT (final concentration was 1% formaldehyde) was added and allowed to incubate under a coverslip for five minutes. The sample was mounted on HistoBond (75x25x1mm from Lamb) slides and freeze-cracked and incubated in methanol at −20°C for one minute and transferred to PBST. Following several washes of PBST the samples were incubated for 30-min in 1% bovine serum albumin diluted in PBST. A hand-cut paraffin square was used to cover the tissue with 50 µL of antibody solution. Incubation was conducted in a humid chamber overnight at 4°C. Slides were rinsed in PBST, then incubated for 2 hours at room temperature with fluorophore-conjugated secondary antibody at a dilution of 1:500. The samples were then mounted in 13 ul of mounting media (20 M N-propyl gallate (Sigma) and 0.14M Tris in glycerol) with a No. 1 ½ (22mm^2^) coverslip and sealed with nail polish.

Primary antibodies were as follows (dilutions are indicated in parentheses): guinea pig anti-ZIM-2 (1:2500; (Phillips and Dernburg 2006), guinea pig anti-PLK-2 (1:750; Harper *et al.* 2011) and rat anti-HIM-8 (1:250; (Phillips and Dernburg 2006) Secondary antibodies were Cy3 anti-rabbit (Jackson Immunochemicals) and Alexa-Fluor 488 anti-guinea pig and anti-rat (Invitrogen).

Quantification of pairing was performed with a minimum of three whole germlines per genotype as in (Phillips *et al.* 2005) on animals 24 hours post L4 stage.

All images were acquired at room temperature using a DeltaVision Personal DV system (Applied Precision) equipped with a 100X N.A. 1.40 oil-immersion objective (Olympus), resulting in an effective XY pixel spacing of 0.064 or 0.040 µm. Images were captured using a “camera” Three-dimensional image stacks were collected at 0.2-µm Z-spacing and processed by constrained, iterative deconvolution. Imaging, image scaling and analysis were performed using functions in the softWoRx software package. Projections were calculated by a maximum intensity algorithm. Composite images were assembled and some false coloring was performed with Adobe Photoshop.

## Acknowledgements

We would like to thank Abby Dernburg and Anne Villeneuve for valuable strains and reagents. The authors declare no competing financial interests. This work was supported by the NIH (grant numbers T32GM008646 [T.B] and R01GM097144 [N.B.]) and the DeAntonio Summer Undergraduate Research Award (G.A.). Some strains were provided by the CGC, which is funded by NIH Office of Research Infrastructure Programs (P40 OD010440).

---

**Figure S1:**
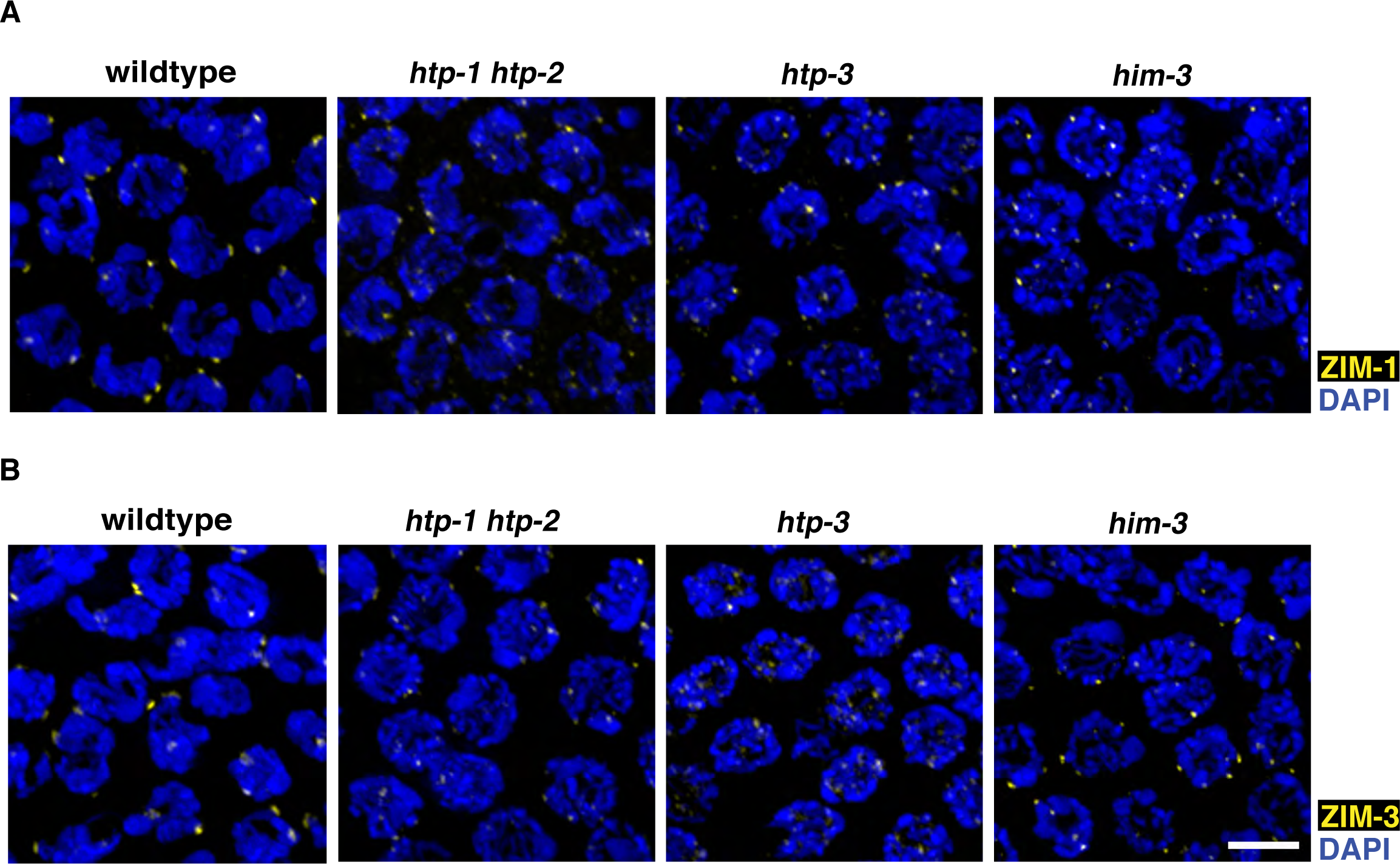
ZIM-1 and ZIM-3 localization in *htp-1 htp-2*, *htp-3* and *him-3* mutants. (A) Images of early meiotic prophase nuclei in wild-type worms, *htp-1/2, htp-3,* and *, him-3* mutants stained to visualize ZIM-1 (yellow) and DAPI (blue). (B) Images of early meiotic prophase nuclei in wild-type worms, *htp-1/2, htp-3,* and *, him-3* mutants stained to visualize ZIM-3 (yellow) and DAPI (blue). Scale bar represents 5 µm.

## References

Aravind, L., and E. V. Koonin, 1998 The HORMA domain: a common structural denominator in mitotic checkpoints, chromosome synapsis and DNA repair. Trends Biochem Sci 23: 284–286.

Bhalla, N., and A. F. Dernburg, 2005 A conserved checkpoint monitors meiotic chromosome synapsis in Caenorhabditis elegans. Science 310: 1683–1686.

Bhalla, N., and A. F. Dernburg, 2008 Prelude to a division. Annu Rev Cell Dev Biol 24: 397–424.

Bhalla, N., D. J. Wynne, V. Jantsch and A. F. Dernburg, 2008 ZHP-3 acts at crossovers to couple meiotic recombination with synaptonemal complex disassembly and bivalent formation in C. elegans. PLoS Genet 4: e1000235.

Bohr, T., C. R. Nelson, E. Klee and N. Bhalla, 2015 Spindle assembly checkpoint proteins regulate and monitor meiotic synapsis in C. elegans. J Cell Biol 211: 233–242.

Brenner, S., 1974 The genetics of Caenorhabditis elegans. Genetics 77: 71–94.

Carballo, J. A., A. L. Johnson, S. G. Sedgwick and R. S. Cha, 2008 Phosphorylation of the axial element protein Hop1 by Mec1/Tel1 ensures meiotic interhomolog recombination. Cell 132: 758–770.

Caryl, A. P., S. J. Armstrong, G. H. Jones and F. C. Franklin, 2000 A homologue of the yeast HOP1 gene is inactivated in the Arabidopsis meiotic mutant asy1. Chromosoma 109: 62–71.

Colaiacovo, M. P., A. J. MacQueen, E. Martinez-Perez, K. Mcdonald, A. Adamo et al., 2003 Synaptonemal complex assembly in C. elegans is dispensable for loading strand-exchange proteins but critical for proper completion of recombination. Dev Cell 5: 463–474.

Couteau, F., K. Nabeshima, A. Villeneuve and M. Zetka, 2004 A component of C. elegans meiotic chromosome axes at the interface of homolog alignment, synapsis, nuclear reorganization, and recombination. Curr Biol 14: 585–592.

Couteau, F., and M. Zetka, 2005 HTP-1 coordinates synaptonemal complex assembly with homolog alignment during meiosis in C. elegans. Genes Dev 19: 2744–2756.

Daniel, K., J. Lange, K. Hached, J. Fu, K. Anastassiadis et al., 2011 Meiotic homologue alignment and its quality surveillance are controlled by mouse HORMAD1. Nat Cell Biol 13: 599–610.

Dernburg, A. F., K. Mcdonald, G. Moulder, R. Barstead, M. Dresser et al., 1998 Meiotic recombination in C. elegans initiates by a conserved mechanism and is dispensable for homologous chromosome synapsis. Cell 94: 387–398.

Derry, W. B., A. P. Putzke and J. H. Rothman, 2001 Caenorhabditis elegans p53: role in apoptosis, meiosis, and stress resistance. Science 294: 591–595.

Deshong, A. J., A. L. Ye, P. Lamelza and N. Bhalla, 2014 A quality control mechanism coordinates meiotic prophase events to promote crossover assurance. PLoS genetics 10: e1004291.

Fukuda, T., K. Daniel, L. Wojtasz, A. Toth and C. Hoog, 2010 A novel mammalian HORMA domain-containing protein, HORMAD1, preferentially associates with unsynapsed meiotic chromosomes. Exp Cell Res 316: 158–171.

Goodyer, W., S. Kaitna, F. Couteau, J. D. Ward, S. J. Boulton et al., 2008 HTP-3 links DSB formation with homolog pairing and crossing over during C. elegans meiosis. Dev Cell 14: 263–274.

Harper, N. C., R. Rillo, S. Jover-Gil, Z. J. Assaf, N. Bhalla et al., 2011 Pairing centers recruit a Polo-like kinase to orchestrate meiotic chromosome dynamics in C. elegans. Developmental cell 21: 934–947.

Hassold, T., and P. Hunt, 2001 To err (meiotically) is human: the genesis of human aneuploidy. Nat Rev Genet 2: 280–291.

Hollingsworth, N. M., L. Goetsch and B. Byers, 1990 The HOP1 gene encodes a meiosis-specific component of yeast chromosomes. Cell 61: 73–84.

Kim, Y., N. Kostow and A. F. Dernburg, 2015 The Chromosome Axis Mediates Feedback Control of CHK-2 to Ensure Crossover Formation in C. elegans. Developmental cell 35: 247–261.

Kim, Y., S. C. Rosenberg, C. L. Kugel, N. Kostow, O. Rog et al., 2014 The chromosome axis controls meiotic events through a hierarchical assembly of HORMA domain proteins. Developmental cell 31: 487–502.

Labella, S., A. Woglar, V. Jantsch and M. Zetka, 2011 Polo kinases establish links between meiotic chromosomes and cytoskeletal forces essential for homolog pairing. Developmental cell 21: 948–958.

MacQueen, A. J., M. P. Colaiacovo, K. Mcdonald and A. M. Villeneuve, 2002 Synapsis-dependent and -independent mechanisms stabilize homolog pairing during meiotic prophase in C. elegans. Genes Dev 16: 2428–2442.

MacQueen, A. J., and A. Hochwagen, 2011 Checkpoint mechanisms: the puppet masters of meiotic prophase. Trends Cell Biol 21: 393–400.

MacQueen, A. J., C. M. Phillips, N. Bhalla, P. Weiser, A. M. Villeneuve et al., 2005 Chromosome Sites Play Dual Roles to Establish Homologous Synapsis during Meiosis in C. elegans. Cell 123: 1037–1050.

Martinez-Perez, E., and A. M. Villeneuve, 2005 HTP-1-dependent constraints coordinate homolog pairing and synapsis and promote chiasma formation during C. elegans meiosis. Genes Dev 19: 2727–2743.

Phillips, C. M., and A. F. Dernburg, 2006 A family of zinc-finger proteins is required for chromosome-specific pairing and synapsis during meiosis in C. elegans. Dev Cell 11: 817–829.

Phillips, C. M., C. Wong, N. Bhalla, P. M. Carlton, P. Weiser et al., 2005 HIM-8 Binds to the X Chromosome Pairing Center and Mediates Chromosome-Specific Meiotic Synapsis. Cell 123: 1051–1063.

Rosu, S., K. A. Zawadzki, E. L. Stamper, D. E. Libuda, A. L. Reese et al., 2013 The C. elegans DSB-2 protein reveals a regulatory network that controls competence for meiotic DSB formation and promotes crossover assurance. PLoS genetics 9: e1003674.

Schumacher, B., K. Hofmann, S. Boulton and A. Gartner, 2001 The C. elegans homolog of the p53 tumor suppressor is required for DNA damage-induced apoptosis. Curr Biol 11: 1722–1727.

Smolikov, S., A. Eizinger, K. Schild-Prufert, A. Hurlburt, K. Mcdonald et al., 2007 SYP-3 restricts synaptonemal complex assembly to bridge paired chromosome axes during meiosis in Caenorhabditis elegans. Genetics 176: 2015–2025.

Smolikov, S., K. Schild-Prufert and M. P. Colaiacovo, 2009 A yeast two-hybrid screen for SYP-3 interactors identifies SYP-4, a component required for synaptonemal complex assembly and chiasma formation in Caenorhabditis elegans meiosis. PLoS Genet 5: e1000669.

Stamper, E. L., S. E. Rodenbusch, S. Rosu, J. Ahringer, A. M. Villeneuve et al., 2013 Identification of DSB-1, a protein required for initiation of meiotic recombination in Caenorhabditis elegans, illuminates a crossover assurance checkpoint. PLoS genetics 9: e1003679.

Woglar, A., A. Daryabeigi, A. Adamo, C. Habacher, T. MacHacek et al., 2013 Matefin/SUN-1 phosphorylation is part of a surveillance mechanism to coordinate chromosome synapsis and recombination with meiotic progression and chromosome movement. PLoS genetics 9: e1003335.

Wojtasz, L., J. M. Cloutier, M. Baumann, K. Daniel, J. Varga et al., 2012 Meiotic DNA double-strand breaks and chromosome asynapsis in mice are monitored by distinct HORMAD2-independent and -dependent mechanisms. Genes & development 26: 958–973.

Wojtasz, L., K. Daniel, I. Roig, E. Bolcun-Filas, H. Xu et al., 2009 Mouse HORMAD1 and HORMAD2, two conserved meiotic chromosomal proteins, are depleted from synapsed chromosome axes with the help of TRIP13 AAA-ATPase. PLoS genetics 5: e1000702.

Zetka, M. C., I. Kawasaki, S. Strome and F. Muller, 1999 Synapsis and chiasma formation in *Caenorhabditis elegans* require HIM-3, a meiotic chromosome core component that functions in chromosome segregation. Genes and Development 13: 2258–2270.

